# Genetic tools for the study of the mangrove killifish*, Kryptolebias marmoratus,* an emerging vertebrate model for phenotypic plasticity

**DOI:** 10.1101/2023.04.21.537589

**Authors:** Cheng-Yu Li, Helena Boldt, Emily Parent, Jax Ficklin, Althea James, Troy J. Anlage, Lena M. Boyer, Brianna R. Pierce, Kellee Siegfried, Matthew P. Harris, Eric S. Haag

## Abstract

*Kryptolebias marmoratus* (Kmar), a teleost fish of the order Cyprinodontiformes, has a suite of unique phenotypes and behaviors not observed in other fishes. Many of these phenotypes are discrete and highly plastic –varying over time within an individual, and in some cases reversible. Kmar and its interfertile sister species, *K. hermaphroditus*, are the only known self-fertile vertebrates. This unusual sexual mode has the potential to provide unique insights into the regulation of vertebrate sexual development, and also lends itself to genetics. Kmar is easily adapted to the lab and requires little maintenance. However, its internal fertilization and small clutch size limits its experimental use. To support Kmar as a genetic model, we compared alternative husbandry techniques to maximize recovery of early cleavage-stage embryos. We find that frequent egg collection enhances yield, and that protease treatment promotes the greatest hatching successes from diapause. We completed a forward mutagenesis screen and recovered several mutant lines that serve as important tools for genetics in this model. Several will serve as useful viable recessive markers for marking crosses. Importantly, the mutant *kissylips* lays embryos at twice the rate of wild-type. Combining frequent egg collection with the *kissylips* mutant background allows for a substantial enhancement of early embryo yield. These improvements were sufficient to allow experimental analysis of early development and the successful mono- and bi-allelic targeted knockout of an endogenous *tyrosinase* gene with CRISPR/Cas9 nucleases. Collectively, these tools will facilitate modern developmental genetics in this fascinating fish, leading to future insights into the regulation of plasticity.

## Introduction

Model organisms, or those species especially amenable to experimentation in a laboratory setting, have been critical in revealing fundaments aspects of developmental biology. Key models such as laboratory mice, Drosophila, the nematode *C. elegans*, and the zebrafish have permitted broad systematic analysis of genetic regulation of development and physiology. However, their characteristics that make them useful experimental genetic models do not capture some important types of regulation observed in nature (Bolker, 1995), and thus cannot be captured in phenotypic screens. One area that has been difficult to study in these workhorse experimental organisms is phenotypic plasticity and its regulation in development and evolution. Notable exceptions include work in nematodes. For example, in *Caenorhabditis elegans* regulation of dauer entry and exit by environmental factors has been extensively studied (Fielenbach and Antebi, 2008). Similarly, genetic screens in *Pristioncius pacificus* have revealed factors comprising a novel signaling network that regulates plasticity in formation of the pharynx, which supports alternative feeding behaviors (Bento et al., 2010; Bui et al., 2018; Sieriebriennikov et al., 2017).

New models that support the study of plasticity should be easily adaptable to the laboratory and provide unique morphological, physiological, or behavioral variants. They should be small, require little care, and have a short generation time. Additionally, ovipary permits easy imaging and manipulation of embryos, and to the one-cell stage bottleneck critical for introduction of transgenes and gene products. An experimental model should also have a genome of manageable size for analysis of variation and mutation.

The mangrove killifish, *Kryptolebias marmoratus* (henceforth “Kmar”; **Figure 1A**), has attracted considerable attention since the pioneering work of Robert W. Harrington, Jr. in the 1960’s (e.g. Harrington, 1961; Harrington and Kallman, 1968; Harrington and Rivas, 1958; Orlando, 2012). Kmar harbors a broad array of unique developmental and physiological adaptations to the extreme environments in which it lives. Some of these traits expose extreme plasticity in their response to environmental change. For example, Kmar can be found out of the water for extended periods (**Figure 1B**). During these emergence times, Kmar epidermis remodels and establishes extensive intra-epithelial capillary networks on the anterior dorsal aspect of its skin (Heffell et al., 2018). Longer periods of emersion lead to reversible gill remodeling (Leblanc et al., 2010; Ong et al., 2007), which is reversed once fish reenter the aqueous environment. Kmar early development also can arrest, having a late embryonic diapause state prior to hatching that resembles a torpor condition of lower metabolic activity (**Figure 1C**). On a shift of environmental conditions these animals will hatch as progress with development and growth.

**Figure 1.**
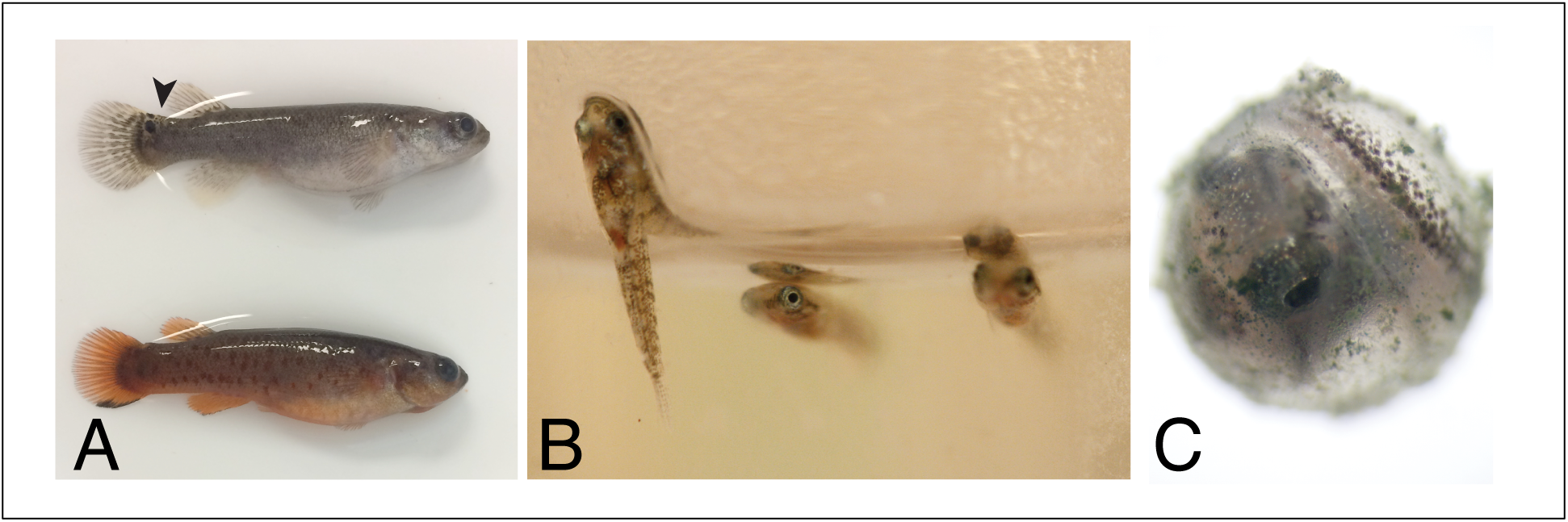
Kmar manifest plastic phenotypes not seen in current vertebrate experimental models of development. A) Kmar reach adulthood as self-fertilizing hermaphrodites (top). Hermaphrodites have a prominent ocellus on tail (arrow head). Males (bottom) are distinguished by orange coloration and a ventral melanic tail bar. B) Emersion behavior and skin respiration is often seen in laboratory culture. C) Pre-hatching developmental diapause.

Most uniquely, Kmar also presents a suite of unusual reproductive adaptations. It is the only known self-fertilizing hermaphrodite vertebrate. Frequent selfing in Kmar has purged most recessive deleterious alleles, allowing completely homozygous strains to be collected from wild populations and maintained without inbreeding depression (Tatarenkov et al., 2012; Tatarenkov et al., 2007; Tatarenkov et al., 2010). Second, Kmar is also a sequential hermaphrodite, with hermaphrodite animals transitioning to become functional males with a frequency and timing that varies between geographical isolates. In both the lab and in nature, Kmar is functionally androdioecious, in that males can fertilize the small proportion of unfertilized eggs released by hermaphrodites (Mackiewicz et al., 2006a; Mackiewicz et al., 2006b; Mackiewicz et al., 2006c). This combination of selfing and outcrossing speeds production of novel homozygous genotypes while still allowing unification of alleles between different strains, much as is done in genetic experimentation in *C. elegans*. These and other phenotypes in Kmar offer an unusually tractable system in which to study plasticity and comparative genetics.

Despite the appealing features of *K. marmoratus*, it differs from medaka and zebrafish in two ways that hinder its use in developmental genetics. First, hermaphrodites rarely lay more than three eggs at a time, unlike the large clutches in zebrafish.. Second, due to internal fertilization only a small fraction of the embryos that are laid are in the zygote or two-cell stage. These attributes hinder the systematic injection and production of morphant, transgenic, or genome-edited strains required to take full advantage of forward genetics. For example, morpholino knockdowns that characterized two mutations impacting Kmar tail development found in a forward screen (Sucar et al., 2016) were conducted with medaka because “it is difficult to obtain many one-cell-stage embryos for … injections” (Saud et al., 2021). Manual coaxing of egg laying can be done for IVF (Nakamura et al., 2008). Further, without intervention a large fraction of *Kmar* embryos never emerge from pre-hatching diapause. To tap the potential of this species for research, it is essential to improve the yield of eggs collected and to increase the survival rates of diapaused embryos.

Here, we describe our work in generating resources and husbandry conditions to improve egg yield and hatching success of *K. marmoratus*. The combination of these assets permitted experimental approaches and production of targeted gene knockouts in the species using CRISPR-Cas9 technology.

## Methods

### Husbandry

Work with *K. marmoratus* in both the Haag and Harris laboratories’ was conducted using protocols approved by the Institutional Animal Care and Use Committees (IACUC) of the University of Maryland, College Park and Boston Children’s Hospital, respectively . Animals were housed in covered, non-circulating tanks, often in isolation from others. For the genetic screen (Harris Lab, Boston MA), fish were maintained in in mouse cages with specialized insets having slots to lets eggs fall to bottom away from fish. These were half filled with zebrafish culture water (sterile, RO with minor salts) amended with 10 parts per thousand (ppt) Instant Ocean salts. Live *Artemia* brine shrimp nauplii were provided ad libitum and allowed to coculture until needing to be replenished (1-2x a week).

For egg-laying, hatching, and CRISPR experiments (Haag Lab, MD), fish were held at 26-28 °C with a 12 hour light, 12 hour dark cycle. Animals were housed singly in one liter polypropylene food containers (Rubbermaid Take-along). Tank lids were tightly closed, with five 2 mm diameter holes drilled near the top to allow gas exchange and feeding via a Pasteur pipette. Each tank contained 750 ml of culture water made with house distilled water and Instant Ocean. Egg-laying and hatching experiments were performed with 25 ppt salt unless otherwise noted. The Haag lab subsequently shifted all fish to 12.5 ppt culture water, and this was used for injections. Adults and juveniles were fed six times a week with approximately 1 ml of a 25% suspension of live *Artemia* nauplii made with distilled water just prior to use. To encourage oviposition, adult hermaphrodites are provided roughly 150 ml of loosely packed polyester fiberfill, either draped in the water by trapping a small portion under the tank lid, or maintained as a floating mass. Eggs deposited on this substrate were gathered with gloved hands, and placed in a 3-inch diameter covered plastic condiment cup containing culture water and held at 26-28 °C. Water was changed every 7-14 days.

### Forward genetic screen

Hon11 is a strain originally collected in the Bay Islands of Honduras (Tatarenkov et al., 2010). Adults were obtained from Brian Ring (Valdosta State Univ., Georgia USA). 16 mature hermaphrodites were mutagenized with 3.0µM N, N, Ethyl-nitrosourea (ENU) once a week for four weeks. Once completed, eggs were collected in batch and collected only once eggs showed consistent viability. Eight hermaphrodites were used as the basis for the screen due to their laying consistent number of progeny (K1-K8). F1 progeny from a single P0 were raised in Petri dishes at 26-29C until hatching, at which point they were transferred in groups of 5-6 to mouse cages on a shelf until maturity. Upon maturity, individual F1 were placed in a mouse cage individually and eggs were collected to screen for phenotypes. Each F1 was given a Roman numeral label to permit phenotype tracking to particular lineages. F2 were collected weekly and were grouped in Petri dishes.

We performed three phenotypic screens:

> <u>Diapause.</u> After a week at room temperature viable eggs are at the start of pre-hatch diapause, and unmutagenized animals do not hatch spontaneously for at least four weeks. F2 fish from mutagenized lines were scored again after two weeks for any batch showing premature hatching at expected Mendelian ratios.
>
> <u>Larval patterning defects.</u> F2 larvae were raised until hatching, and morphology was scored prior to moving to larger tanks.
>
> <u>Adult phenotypes</u>. F2 progeny were raised until mature at which point outward morphology and reproductive phenotypes were scored. Reproductive phenotypes screened included sex ratio and sterility. A large proportion of F2 progeny were scored for regeneration potential of the caudal fin.

### Egg Laying Experiments

Egg counts for weekly collections were established for each fish over two months. For twice daily treatments, each experiment began by clearing the fiberfill mop of all eggs that may have been present. Eggs were subsequently collected shortly after the lights coming on at 8am and again at 7pm (i.e. just before the lights went off), repeated for 5-7 days. Fish that failed to lay any embryos were not included in analysis. One SOB10 fish in the “teabag experiment” (Figure 4B) produced 16 eggs in one 10 hour period, an extreme outlier that was also excluded.

### Diapaused embryo hatching success

Embryos that received control conditions were reared using the conditions previously described, with a water change approximately two weeks after oviposition. By this age, all viable embryos have completed development and are in a pre-hatch diapause state. For embryos in the “Reduced Salinity” treatment group, the first water change was with half-strength (12.5) ppt salinity water. Embryos in the “Agitation” treatment group received 25 ppt salinity water but were placed on an orbital platform shaker set at 60 rpm. Two Pronase treatments were initially used: Protocol 1) 0.02% pronase in 25 ppt water, held without a water change, and Protocol 2) .05% pronase in 25 ppt water for two days, followed by a change to 25 ppt water lacking Pronase.

The higher concentration of Pronase effectively stimulated hatching, but was accompanied by high rates of perinatal larval death. The data reported in **Figure 5** were all from Protocol 1.

### CRISPR-Cas9 mutagenesis

We modelled our targeting of the *Tyrosinase* gene on similar work in the cichlid *Astatotilapia burtoni* (Li et al., 2021).

#### Design and synthesis of guide RNAs

The *K. marmoratus Tyrosinase* gene sequence from NCBI Genbank (Gene ID: 108244618, NCBI Reference Sequence: NC_051431.1). The IDT Custom Alt-R™ CRISPR-Cas9 guide RNA online tool (https://www.idtdna.com/site/order/ <u>designtool/index/CRISPR_SEQUENCE</u>) was used to design guide RNA (gRNA) targeting exon 4. Cleavage within this exon has successfully abolished melanocyte formation in *A. burtoni* (Li et al., 2021). We chose two gRNA targeting sites 51 bp apart (Figure 1A), to allow for double cut-mediated deletions while still preserving overall mRNA splicing splicing. gRNA sequences were used as BLAST queries (Altschul et al., 1990) against the *K. marmoratus* genome to confirm absence of obvious off-target effects. Duplexed synthetic trans-activating CRISPR RNA (tracrRNA) and target-specific CRISPR RNA (crRNA) (both from IDT, Coralville, IA) were used instead of single guide RNA (sgRNA), as this improves editing rates in zebrafish (Hoshijima et al., 2019).

#### Injection cocktail

crRNA and tracrRNA were first individually diluted as 100 μM stock solutions in manufacturer-provided duplex buffer (IDT; 30 mM Hepes pH 7.5, 100 mM Potassium acetate). To create crRNA:tracrRNA duplexes, equal volumes of crRNA and tracrRNA stock solutions were mixed and annealed in a thermocycler using this program: 95°C, 5 min; cool at 0.1°C/sec to 25°C; 25°C, 5 min; cool to 4°C rapidly. The 50 μM crRNA:tracrRNA duplex stock was further diluted 1:1 with duplex buffer to produce a 25 μM working stock. Cas9 protein (IDT Alt-R S.p. Cas9 Nuclease V3, 1081058) was adjusted to 25 μM stock solution in 20 mM HEPES-NaOH (pH 7.5), 350 mM KCl, 20% glycerol. Injection mixtures combined the following: 5 μM Cas9 protein; 5 μM crRNA:tracrRNA duplex; 0.25% Texas Red conjugated dextran (final conc. 0.25%).

##### Microinjection of embryos

On the day of the injection, we checked and collected eggs from individually house *kissy lips* tanks twice: the first collection in the morning between 08:30-09:00 (shortly after lights on); the second one was between 12:00-12:30. To increase the genome-editing rate, only embryos between 1- to 16-cell stage (Figure 1B) were injected. We performed five rounds of injections in a total of 77 such early cleavage stage embryos.

Prior to microinjection, the gRNA:Cas9 ribonucleoprotein complex solution was incubated at 37°C for 5 min and then backfilled into 3 microinjection needles using an Eppendorf GELoader tip (Eppendorf, Cat# 022351656). After loading embryos into the embryo holder which covered with 12.5 ppt of salinity water with 0.0001% of methylene blue, we injected them with 4 pulses of 1 nL, using 2.5 ms pulse duration at 22 psi. To further explore the editing efficiency in different conditions, 53 eggs were directly injected into blastomere, and 24 eggs were injected into yolk (Figure 1B). Once injections were complete, embryos were gently removed from trenches using a micro laboratory spatula (Fisher, blade size: 19 mm × 4.8 mm) and then transferred individually into 12-well plates with 2 mL of 12.5 PPT water containing 0.0001% of methylene blue. Embryo survival and pigmentation were monitored for approximately 28 days post-injection.

Microinjections were performed under a stereomicroscope (Nikon, SMZ745) equipped with a 3D micromanipulator (Narishige, M-152). Injection pulses were driven by a tank of compressed dry air (size 200, adapter CGA-590) fed into a Milli-Pulse Pressure Injector (Applied Scientific Instrumentation; MPPI-3). Embryos were immobilized for injection in grooves cast in a 2.0% agarose block prepared with 12.5 ppt salinity water. Grooves were formed by addition of several parallel Micropet Disposable Pipettes (Clay Adams, ACCU-FILL90, 20µL, #4618) in a standard Petri plate. Removal of the micropipettes after gelling produces 1.5 mm-wide trenches (Figure 1C). Microinjection needles was made from glass capillary tubes (GC100F-10, 1.0 mm O.D; 0.58 mm I.D; Harvard Apparatus) pulled with a Sutter P-97 micropipette needle puller. Initially sealed needle tips were broken by holding taut a Kimwipe and gently tapping the tip straight into it. Needles with the ideal needle bore size of 10-12.5 µm outer diameter were identified using a microscope equipped with an ocular micrometer. This diameter of needle delivers ∼1 nL of solution when a 2.5 ms air pulse at 22 p.s.i.

##### Indel mutation quantification

PCR and Sanger sequencing were used to assess genome-editing efficiency in each CRISPR-injected G0 fish. We designed primer pairs (**Table 1**) to amplify ∼300bp region surrounding the Exon 4 target sites (**Figure 1A**). At 3 months of age, we collected about ∼1 mm^2^ of tailfin tissue from injected fish to identify those carrying the high rates of mutation. These fish with somatic mutations are preferred for breeding, as they are most likely to also carry mutations in the germline. Genomic DNA was extracted using a modified HotSHOT approach (Truett et al., 2000). The fin-clipped samples were first transferred into 0.2 mL PCR tubes, followed by adding 180 µL of NaOH (50 mM). The samples were then heated at 95°C for 15 min and cool to 4°C. Lastly, 20 µL of Tris buffer (1 M, pH 8.0) was added to each sample to neutralize pH.

**Table 1.** Summary of forward screen phenotypes.

| **Affected Phenotype** | **# Mutants Identified/**<br>**# Genomes** | **# Re-identified** | **Designation of retained mutant** |
| --- | --- | --- | --- |
| **Larval morphology** | 81/602 | 2 | *Albino, stumpy* |
| **Diapause** | 28/602 | 6 |  |
| **Larval total** | 109/602; 0.18 |  |  |
| **Adult morphology** | 91/602 | 1 | *kissy lips* |
| **Regeneration** | 0/512 |  |  |
| **Sex determination** | 2/602 | 2 | *mher-1, mher-2* |
| **Fertility** | 11/392 | - |  |
| **total** | 104/602; 0.17 |  |  |

PCR primers used for genotyping: 5’TGTAAAACGACGGCCAGTtgctgtcgtttctgctcctgtg 3’ [capitalized portion of the primer sequence is an M13 universal tag, permitting subsequent reamplification with a fluorescent dye-conjugated M13 primer for fragment analysis]. Reverse primer 5’ agcgagattgcaagcgtattgc 3’. 35 cycles of amplification were performed with 94°C, 2 min; 94°C, 15 s, 60°C, 15 s, 72°C, 30 s; followed by a polishing step of 72°C, 7 min. 2% agarose gels were run to confirm successful amplification. Most indels are <15 bp in size and cannot be resolved by this method, so amplicons were Sanger sequenced. The indel mutation rates were analyzed by using the web tool Synthego ICE (v3.0. https://ice.synthego.com/) and TIDE (Tracking of Indels by Decomposition; Brinkman et al., 2014)).

To assess if the mutations were transmitted into the germline cells and passing to the offspring, we also collected >50 eggs from each surviving P0 injected fish, and screened for pigment loss in the F1 generation.

## Results

### Chemical mutagenesis screen to identify markers for genetic analyses and analysis tools

To assess feasibility of forward genetic screens in Kmar as well as to establish tools to support genetic analyses, we initiated a chemical mutagenesis in the Hon11 wild-type background.

Hon11 was previously identified as being hearty and a consistent egg producer in the lab (Sucar et al., 2016). We mutagenized 11 Hon11 hermaphrodites 4 times with 3.0µM ENU, using protocol identical to that used for zebrafish (Rohner et al., 2011). Unlike a prior screen in Kmar (Sucar et al., 2016), we did not collect eggs in batch from selfing groups, but rather separated individual P0 founders and collected eggs daily, raising F1 of each founder to track potential clonal mutations in individual germlines. In this genetic screen design (**Figure 2**) we could more precisely associate expressivity of particular phenotypes as well as assess frequency of phenotypes within a particular founder lineages.

**Figure 2.**
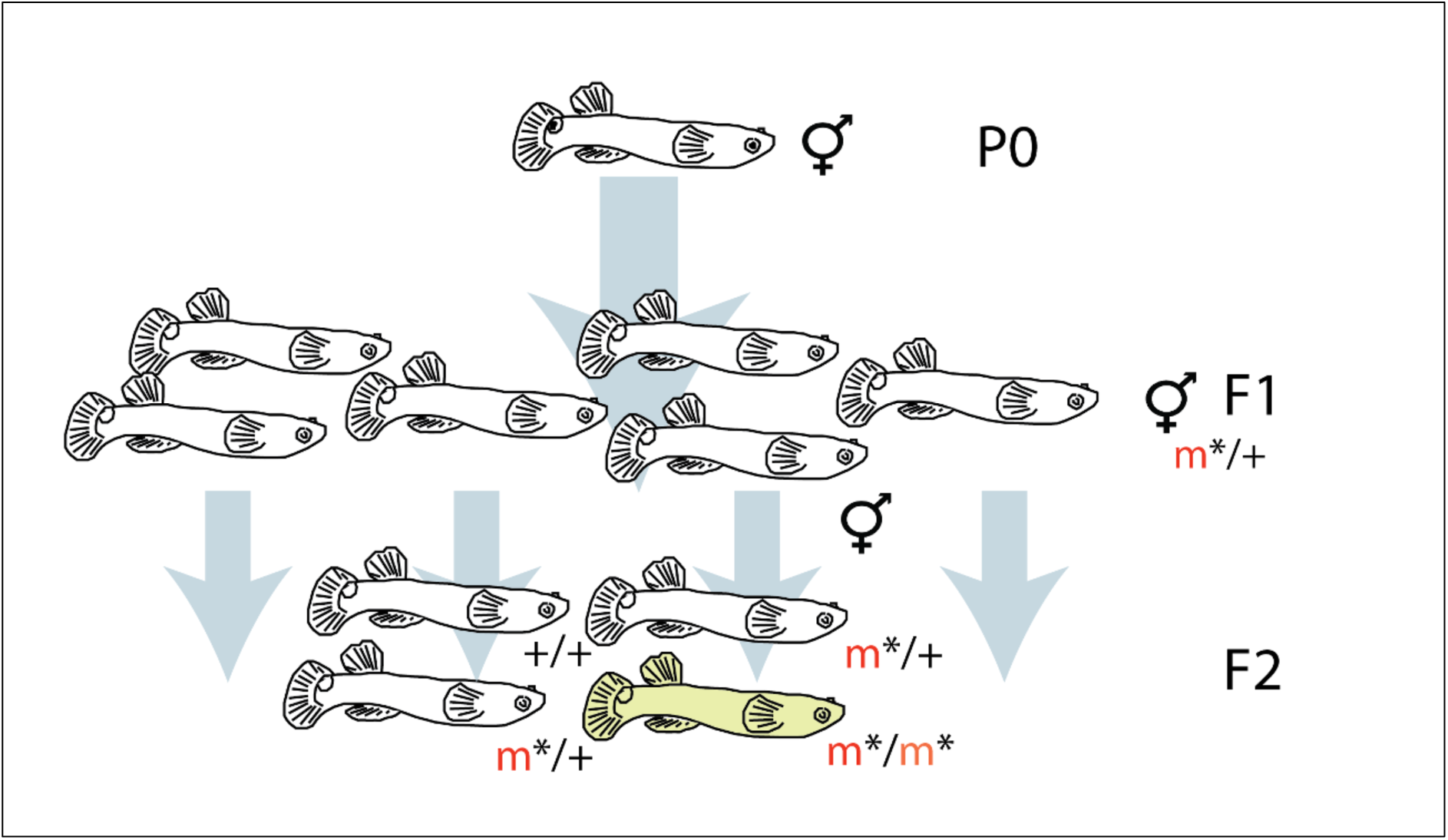
Design of mutagenesis screen in Kmar. Adult hermaphrodite parents (P0) were mutagenized and allowed to lay F1 self-progeny, which are often heterozygous for newly induced recessive mutations (m*/+). Selfing of these founders produces roughly 25% homozygous F2 progeny (m*/m*), which manifest a range of phenotypes. This path to novel homozygous mutants requires far fewer animals and one fewer generation than similar screens in obligately outcrossing species.

In sum >450 F1 founders were produced. Progeny of 301 founders were screened for mutations affecting entry or maintenance of diapause, larval and adult morphologies. Of these, 261 F1 progeny were screened for regeneration and 196 for fertility phenotypes (**Table 1**). No F1 (presumably dominant) phenotypes were observed in the screen, nor were dominant mutations observed in the F2. We identified a large spectrum of mutants in the F2, many of which were subsequently reidentified in crosses of isolated homozygous founders, or in carriers in the case of larval lethal or adult sterile phenotypes (**Table 1**). Mutants were identified in most classes analyzed, however we did not detect any that affected tail fin regeneration. Although Kmar has distinct background pigmentation and has a prominent ocellus on the dorsoposterior caudal peduncle, only one pigmentation mutant was identified, resulting in a classic *albino* phenotype. Due to the number of mutants identified, re-identification efforts were performed only on a subset of these mutants having the most reproducible and useful phenotypes (see below).

Unlike zebrafish and medaka screens, in which mutagenized P0 and F1 are outcrossed, our Kmar screen selfed mutagenized founders. As a result, mutation burden is maintained in the population. Replicated phenotypes suggestive of a generalized response to mutational burden were mainly seen in mutants affecting early development. Such mutants often had short embryonic axes or lack of eyes. Some survived to be axially deformed or eyeless adults, respectively. These were not maintained.

#### Increased tolerance for high-density rearing

*K. marmoratus* live as a solitary animals in confined spaces within mangrove forests (Taylor, 2012). In the lab, most lines individuals show high level of agression when housed in close quarters with conspecifics. This usually leads to lower fecundity and sometimes death. As it relates to lab husbandry and screens, such antagonistic behavior is a detriment to optimizing housing and increases space requirements for fishes.

In the breeding strategy as part of the screen individuals were housed in moderate densities that increased over time as the number of unique populations were refined in the F2 mutants and more individuals with comparable genotypes were raised. Addition of fiberfill and small vinyl tubing permitted local environments and barriers that helped increase rearing densities. Kmar raised in such manner became, or were selected to be, tolerant of communal living, being able to be kept up 10-12 per 500ml without conflict. As the lines were initially isogenic, the changes in these lines were presumably due to acclimation rather than selection of particular background or induced variants. A byproduct of this acclimation is that in most popoulations, the size of reproductive adults is considerably smaller than singly housed adults of the founder strains. We have not tested this specifically, but it is likely that once placed in isolation these fish will continue growth as most teleosts have indeterminant growth through their lifespan.

### Genetic lines maintained for use as genetic markers

Five healthy mutant lines with interesting phenotypes were retained for further characterization:

> *Stumpy:* These mutants have a shortened body, but are otherwise normal hermaphrodites.
>
> *Male hermaphrodite (mher-1, mher-2):* These two lines initially produced a high frequency of male F2 progeny, suggestive of a segregating sex determination defect. Over time, however, excess male production waned, and microsatellite-verified mapping crosses between *mher* males and hermaphrodites of the SOB10 wild-type strain failed to produce F2 males (not shown). These lines were abandoned, but their existence indicates that sexual plasticity can be impacted by mutagenesis.
>
> *Albino: Albino* mutants lack pigment, a defect that can be observed very early in larval development (**Figure 3A,B**). Such mutants are of great utility in visualization of development and localization of gene expression by in situ hybridization or transgenesis. In addition, crosses between *albino* hermaphrodite founders and pigmented males allow early recognition of outcross progeny without more complex genotyping.
>
> *Kissylips:* Fish with this mutation have a general craniofacial shape change and may have postaxial skeletal phenotype, however not dysmorphic or pathological. *kissylips* gets its name from the novel pattering of a stripe of irridiophores across the distal jaw resembling the appearance of lipstick (**Figure 3C,D**). We fortuitously discovered that this mutant line is super-fecund and the line was set aside for continued analysis (**Figure 4**).

**Figure 3.**
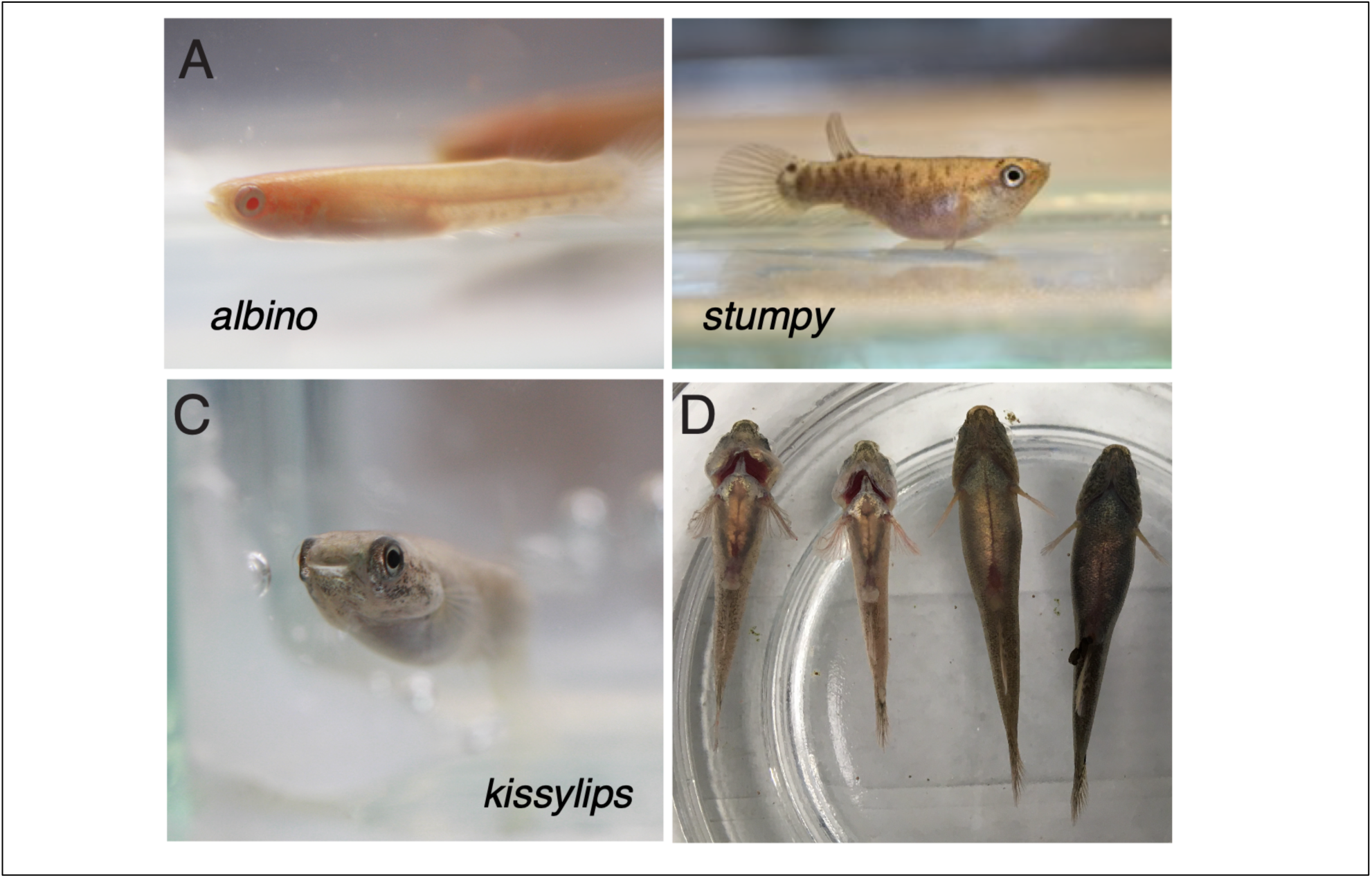
Forward genetic mutant lines bolstering Kmar experimental utility. A) Adult *albino* mutant Kmar show lack of melanin differentiation both in body and retinal pigmentation. No other phenotypes are observable. B. *stumpy* mutants are compressed along their rostral-caudal axis. C) *kissylips* mutant adults have a irridiophore stripe along the distal jaws D) At 8 months of age, *kissylips* (two at left) are smaller than 5-6 month-old wild-type fish (two at right), and have constitutively flared opercula. *kissylips* is also hyperfecund (Figure 4).

**Figure 4.**
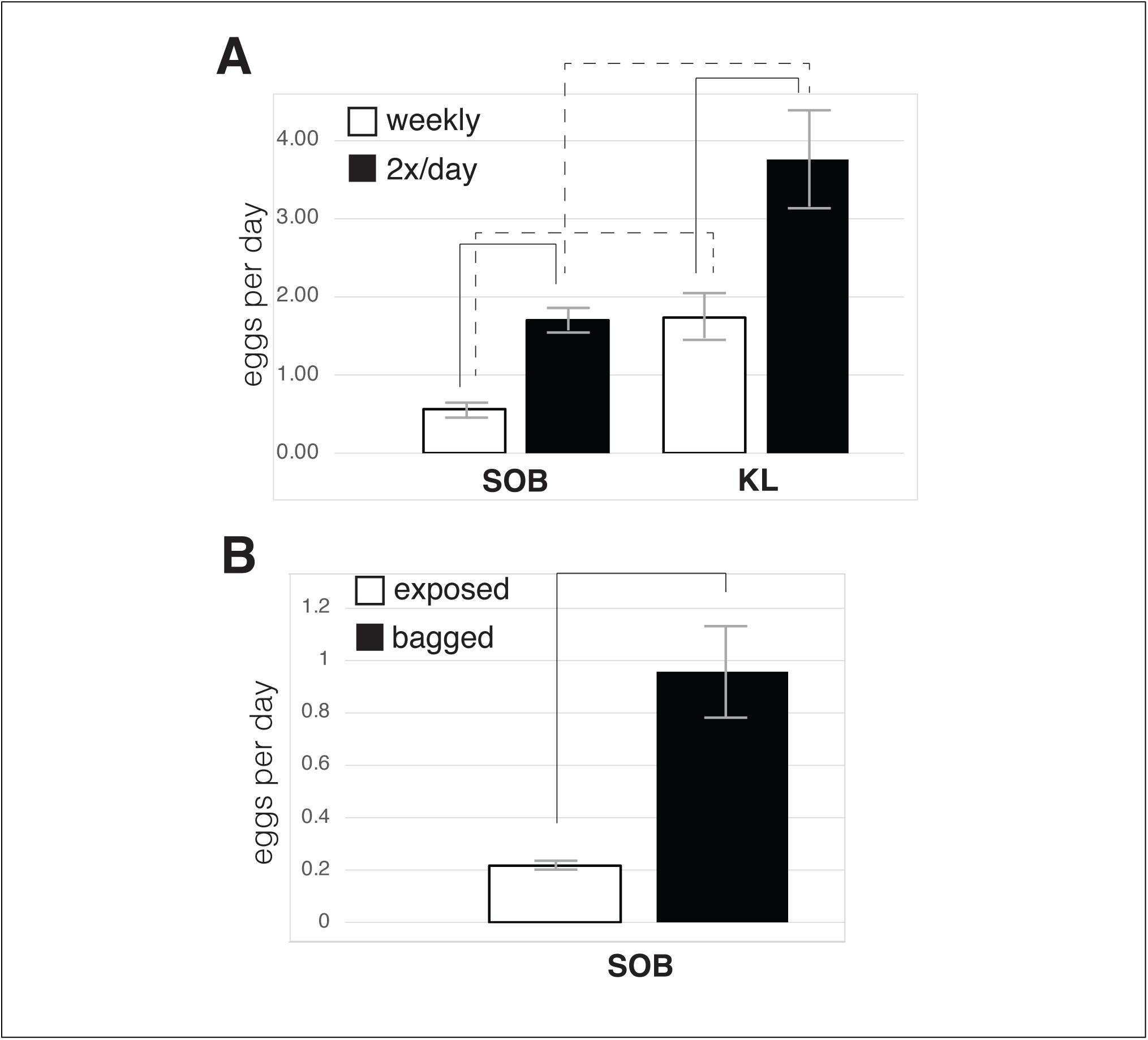
Factors influencing egg yield. A. Frequency of egg removal. Eggs were collected from wild-type SOB8 and SOB strains (collectively “SOB”, N=10) and the HON11-derived mutant *kissy lips* line (“KL,” N=7)) weekly, and then twice per day. Solid lines indicate significant increases in egg yields within a group (paired sign test, *P<*0.01). In both collection schemes, *kissy lips* yield significantly more eggs than SOB8/10, as indicated by dashed lines (two-tailed T-test, *P<*0.01). B. Protection of eggs. Eggs from SOB8/10 hermaphrodites (N=14) were either collected weekly as in A (“exposed”), or transferred up to twice per day into paper teabags and left in the tanks to accumulate (“bagged”). Bagging of eggs significantly increased yield, as indicated by the solid line (two-tailed T-test, *P<*0.01).

### Husbandry factors impacting egg yield

Harrington (1963) reported that for his Floridian strains, hermaphrodites lay eggs at all hours of the day, but peak mid-day. To understand the laying patterns of our various strains of *K. marmoratus*, we monitored the number of eggs collected from hermaphrodites of the inbred wild-type lines SOB8 and SOB10, and also for the Hon11-derived *kissylips* mutant. After two months of weekly collections, we began collecting eggs from these same fish twice a day, once within two hours of lights-on (8am) in the morning, and again within two hours of lights off (8pm). We did not find any qualitative difference in yield between collection times (not shown), indicating that these three strains do not have a strong preference for mid-day laying. However, we did observe that the twice daily collection roughly doubled the yield of eggs per day over the weekly collections in both wild-type and *kissy-lips* strains (**Figure 4A**). Yield from *kissy lips* mutants were roughly double SOB strains in both collection regimes, such that frequent collection of from the mutants quadruples the egg yield compared to weekly collection of wild-type stocks.

The increased yield seen with more frequent collections might be due to the stimulation of oviposition. For example, the presence of an egg in the spawning mop may repress egg laying via a chemosensory mechanism. However, as is common in zebrafish, Harrington (1963) observed that hermaphrodites often eat their eggs soon after oviposition. Thus, the improved yield from frequent collections may simply indicate that *K. marmoratus* eat a greater fraction of their eggs when left for long periods in their tanks. To distinguish between these hypotheses, we repeated the twice daily collections for wild-type fish, but instead of removing the eggs they were instead placed inside unbleached paper tea bags, and then returned the tank. We reasoned that this would allow fish to sense and possibly interact with eggs without the ability to eat them. With eggs accumulating in the tea bags, the enhanced yield in the twice-daily collections remained (**Figure 4B**).

### Factors influencing larval hatching

*K. marmoratus* embryos undergo pre-hatching larval diapause. However, in our lab colonies many otherwise viable embryos fail to hatch into larvae, a phenomenon dubbed “stubborn eggs” (Turner et al., 2006). We tested two factors that had been suggested to increase hatching, abrupt introduction of lower-salinity culture water (from 25 PPT to 10 PPT) and agitation of embryos using a rotary shaker. We also tested the fungal protease cocktail Pronase, which has been reported to stimulate hatching (Kanamori et al. 2006). We found that agitation had no impact on hatching, but that reduction of salinity did increase hatching somewhat (**Figure 5**). However, Pronase had a much larger effect. Since these controlled experiments were performed, Pronase treatment has become our standard protocol to maximize hatching. It is common to obtain over 90% success if treatment is initiated at 6-8 weeks after oviposition and juveniles are removed from treated water shortly after hatching.

**Figure 5.**
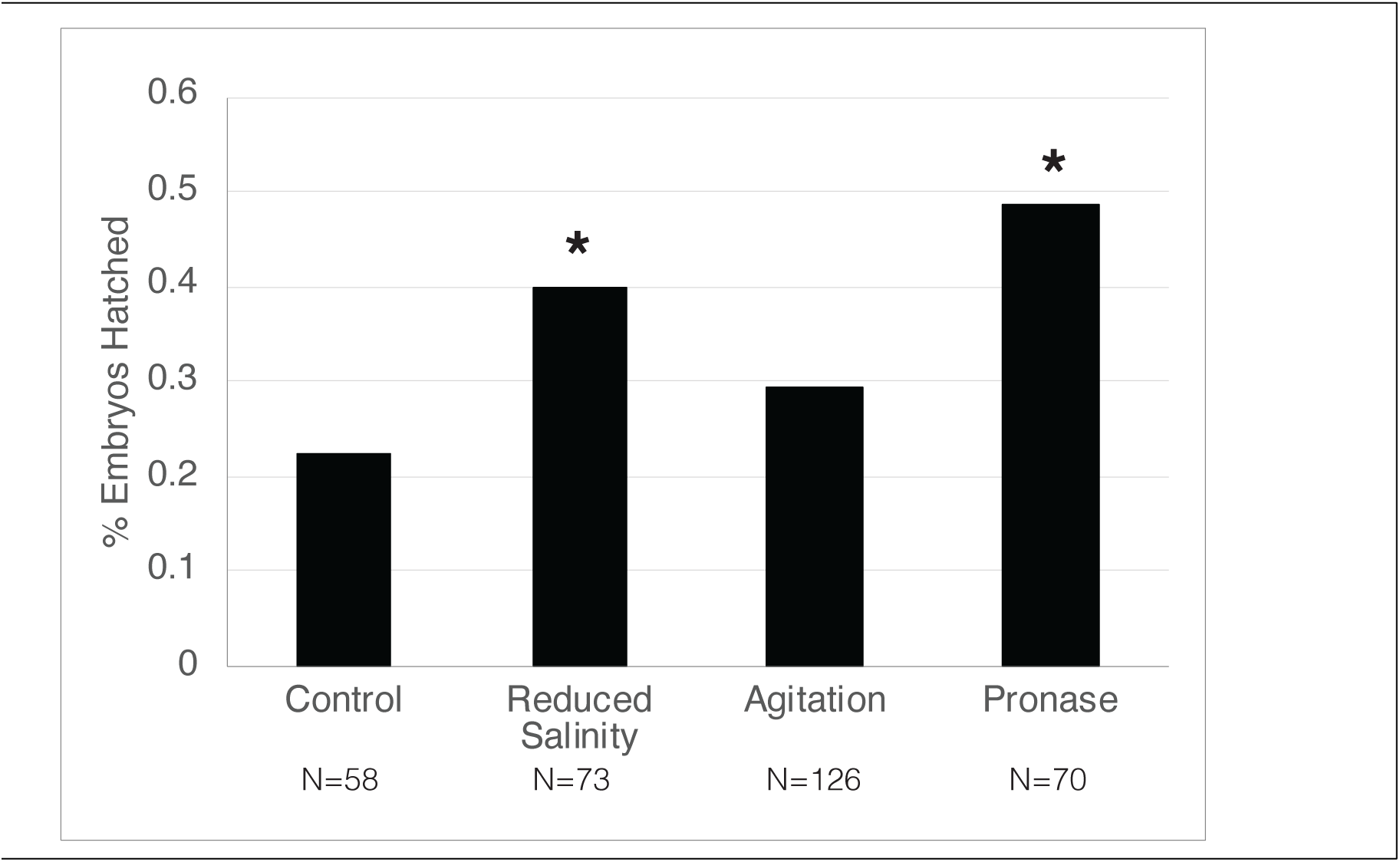
Impacts of various treatments on hatching of diapaused embryos. Compared to control embryos left in standard culture water without agitation, diapaused embryos that were shifted to reduced salinity or to culture water amended with pronase hatched at a significantly higher rate (as indicted by asterisks, chi-squared test, 1 d.f. P<.001). Agitation, in contrast, did not significantly impact hatching success.

### CRISPR/Cas9 gene disruption

A general strategy for injection of *K. marmoratus* embryos has been described (Mourabit et al., 2011; Mourabit and Kudoh, 2012). By combining a large colony of *kissylips* mutants with frequent egg collections, enough embryos in the 1-4-cell stages of development can be obtained to allow daily injections (**Figure 6**). We used this to achieve a targeted knockout.

**Figure 6.**
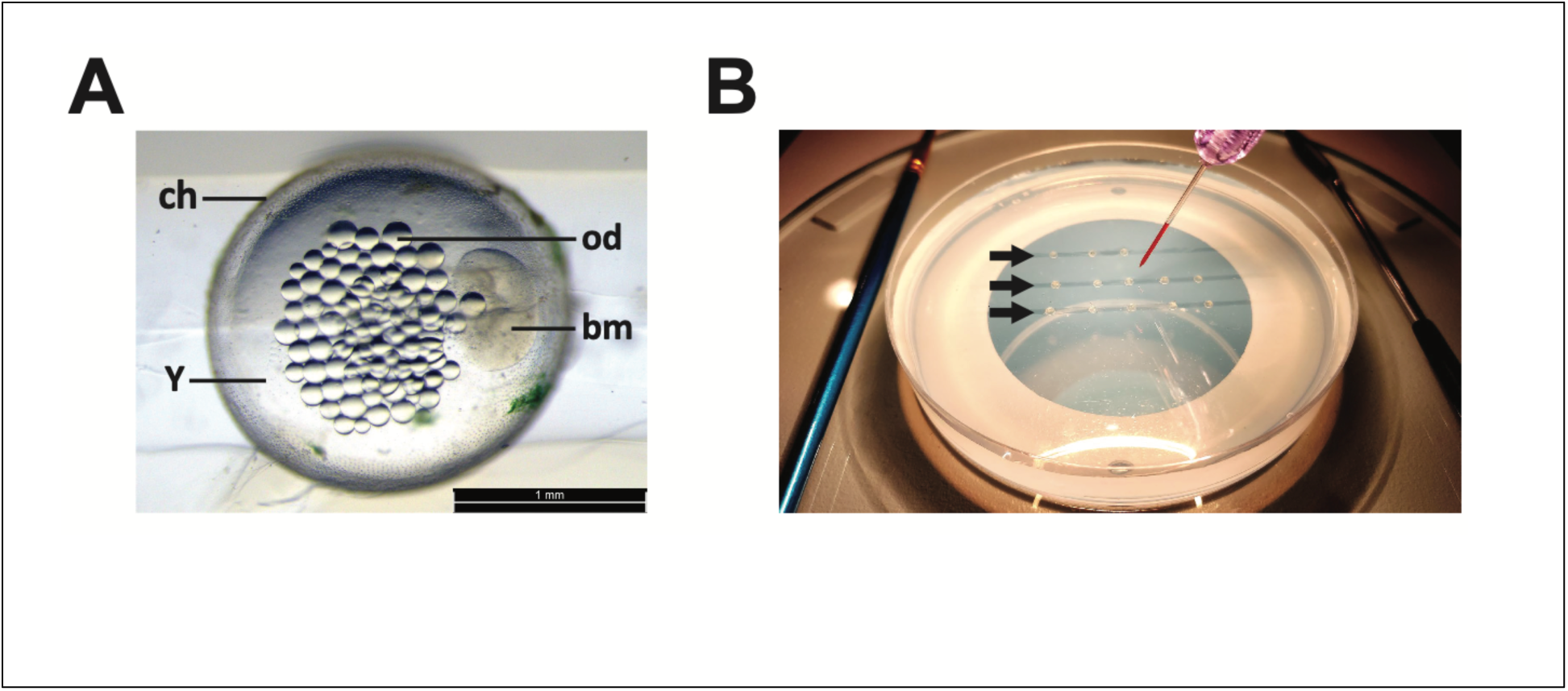
Injection of *K. marmoratus* embryos. A. A representative two-cell *K. marmoratus* embryo, showing the blastomeres (bm), chorion (ch), oil droplets (od), and yolk (y). C. Injection configuration, on the stage of a stereomicroscope. Black arrows indicate grooves cast into an agarose plate, with embryos lined up in each. To the left is the small spatula used for inserting and retrieving embryos, and at upper right the needle and a dye-filled needle are visible.

We injected 77 eggs with CRISPR/Cas9 nucleases targeting the *tyrosinase* gene; 53 directly into blastomeres, and 24 eggs into yolk. 18 injected embryos survived to adulthood, but survival after blastomere injection was half of that of yolk injection (yolk injection: 9/24 survived; blastomere injection: 9/53 survived; **Figure 7A**). However, blastomere injection significantly increased the editing efficiency in embryos that did survive (*Mann-Whitney* test, df=1, *P* = 0.0023; **Figure 7B**). One P0 CRISPR-injected embryo grew to be a complete albino larva, while another had only sparse pigment (**Figure 7C**). This indicates CRISPR-Cas9 editing occurred to a variable extent in P0 injectees, with biallelic *tyrosinase* knockouts in some or all melanocytes. Sequencing analysis confirmed the existence of indel mutations at one of the two target sites (**Figure 7D, Table 2**). Synthego ICE analysis further revealed that these mutant animals carried multiple mutant alleles (**Figure 7E**). To identify pigmented carriers, pigmented progeny of P0 injectees were screened for pigment-loss phenotypes. Two produced multiple albino F1 offspring (**Figure 7F, Table 2**). The fraction of albinos, between 4%-15%, is significantly less than the 25% expected for a homogenous heterozygote, indicating that these individuals have mosaic germ cells, with some wild-type and some bearing monoallelic mutations (**Figure 7E**).

**Figure 7.**
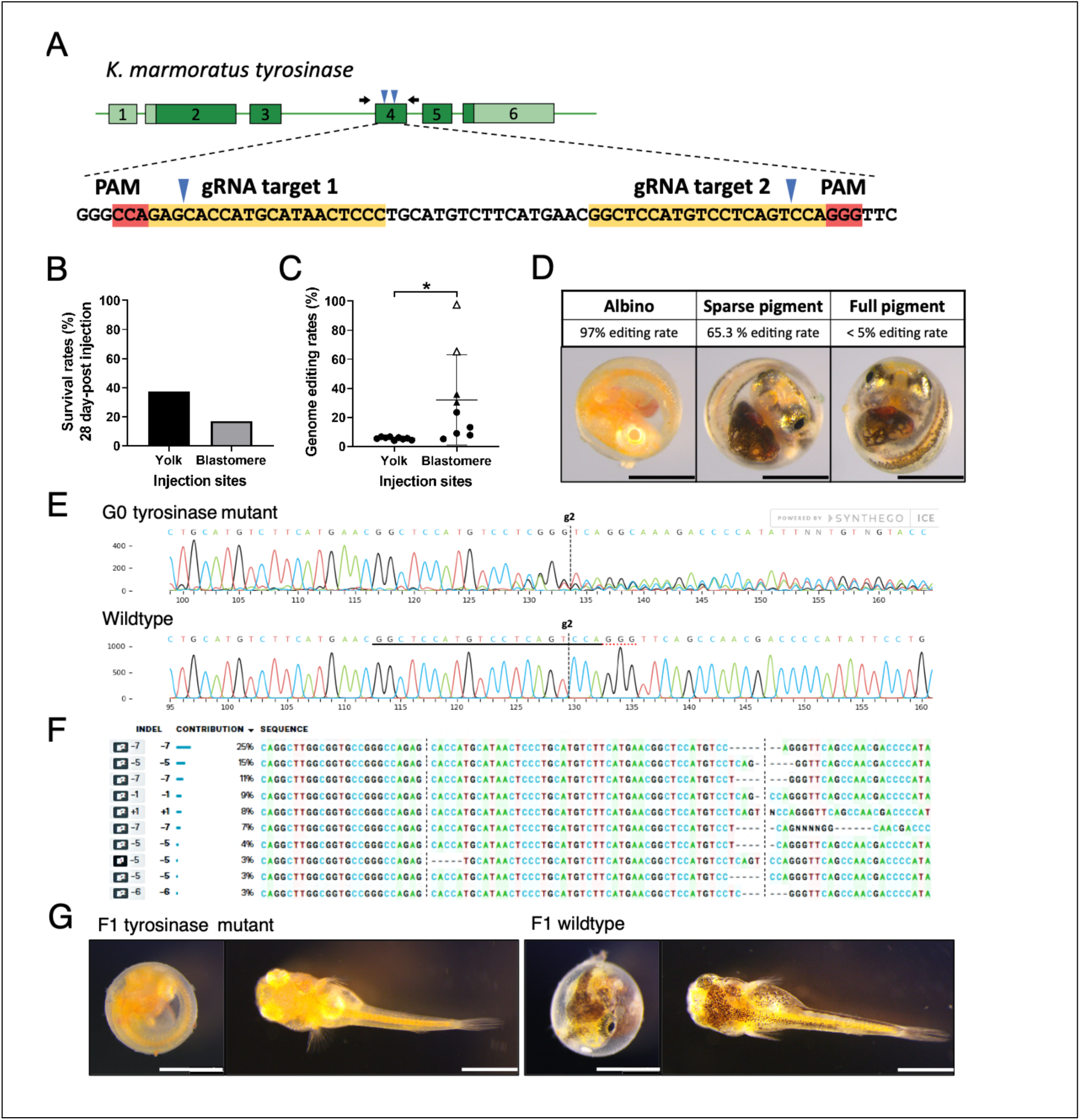
Targeted disruption of *tyrosinase* gene by CRISPR/Cas9 mutagenesis. A. Structure of the *K. marmoratus tyrosinase* gene, with boxes representing exons and darker shading coding sequences. Blue carets depict the locations of target sequences for guide RNAs, with the sequence (target 2) or the sequence’s complement (target 1) highlighted in yellow below. Black arrows indicate the primers used for PCR amplification of the target region. Survival rates (B) in embryos receiving CRISPR injection from blastomere (N=53) or yolk (N=24). C. Genome editing rates in the 18 surviving fish, as measured by target amplicon sequencing. Open triangles represent the two embryos that had severe pigment-loss phenotype; black triangles represent the two individuals producing F1 albino progeny. D. Three representative phenotypes and the corresponding gene editing rates in G0 CRISPR-injected embryos. E. Sanger sequencing data of an exon 4 amplicon from a G0 albino that died shortly after hatching, aligned with the gene sequence around the gRNA target 2. Abrupt sequence heterogeneity beginning near the cut site suggests a mixture of mutations were induced in this sample. (E) Synthego ICE analysis showed all the possible insertions/deletions (indels) in this G0 albino mutant. (F) Sibling F1 embryos from a normally pigmented G0 progenitor, showing normal (right) and melanin-free mutants (left), both in pre-hatching diapause and after hatching.

**Table 2.** Gene editing rates in P0 CRISPR-injected individuals and F1 mutant recovery from P0 crispant.

| Embryo ID | Injection sites | Phenotype | P0 Editing rates | F1 albino/total | F1 albino % | Note |
| --- | --- | --- | --- | --- | --- | --- |
| 8 | Blastomere | wildtype | **30.40%** | **3/78** | **3.85%** | ~39% mutant gametocytes |
| 13 | Blastomere | wildtype | 9.10% | 0/60 | 0.00% | mutant cells only in soma |
| 16 | Blastomere | wildtype | 23.40% | 0/89 | 0.00% | mutant cells only in soma |
| 17 | Blastomere | wildtype | 7.70% | N/A | N/A | did not produce eggs |
| 19 | Yolk | wildtype | 6.60% | 0/84 | 0.00% | mutant cells only in soma |
| 22 | Yolk | wildtype | 5.60% | 0/52 | 0.00% | mutant cells only in soma |
| 29 | Blastomere | wildtype | 5.10% | 0/91 | 0.00% | mutant cells only in soma |
| 32 | Blastomere | wildtype | **35.80%** | **17/129** | **13.18%** | ~73% mutant gametocytes |
| 34 | Yolk | wildtype | 5% | 0/119 | 0.00% | mutant cells only in soma |
| 37 | Yolk | wildtype | 6.10% | 0/91 | 0.00% | mutant cells only in soma |
| 40 | Yolk | wildtype | 6.60% | 0/140 | 0.00% | mutant cells only in soma |
| 43 | Yolk | wildtype | 4.30% | 0/143 | 0.00% | mutant cells only in soma |
| 44 | Blastomere | **Sparse pigment** | **65.30%** | N/A | N/A | died as larva |
| 47 | Yolk | wildtype | 5.90% | 0/110 | 0.00% | mutant cells only in soma |
| 52 | Blastomere | **Albino** | **97.20%** | N/A | N/A | died as larva |
| 59 | Yolk | wildtype | 4.10% | 0/13 | 0.00% | mutant cells only in soma |
| 60 | Yolk | wildtype | 5.40% | N/A | N/A | died as larva |
| 66 | Blastomere | wildtype | 13.20% | 0/32 | 0.00% | mutant cells only in soma |

## Discussion

### Comparison of efficiency in genetic screens among small laboratory fish models

Despite the advent of targeted gene editing in a growing range of organisms (Reardon, 2019), unbiased forward genetic screens still remain a powerful tool for the genetic analysis of phenotype. The execution of large-scale forward genetic screens in the zebrafish, and complemented with similar designs in the medaka, proved the utility of such approaches in a vertebrate model, as had been previously shown in invertebrates such as Drosophila and *C. elegans*. One of the demands on doing such screens is the number of individuals needed to identify novel homozygous recessive mutants. In obligate outcrossing animals, one must carry analysis into the F3 progeny of isolated F2 families, only ¼ of which will have two mutation-carrying parents. A strength of *C. elegans* is the use of selfing within screen designs to reveal new recessives in F2 progeny of single F1 founders. Kmar’s unique sexual strategy provides a similar approach (**Figure 2**).

We successfully screened mutagenized Kmar for recessive alleles affecting a varied array of larval and adult phenotypes. Since circulating water was not necessary, the work here was completed on only one tank rack, or shelf space. Additionally, we only used about 1/300^th^ of the fish that would have been necessary in a comparable zebrafish screen. Unlike prior screens described in Kmar (Moore et al., 2012; Sucar et al., 2016), we kept individual lines isolated to assess for lineage-bias in alleles and potential mitotic clonal mutations arising. We isolated a wide array of mutant phenotypes. No apparent bias in mutation type with founder was seen. Overall mutagenesis efficiency was comparable to that seen in other screens (**Table 3**), with 36 recessive mutants observed in larvae and 81 in adults. These data confirm that chemical mutagenesis screens are a viable experimental approach in Kmar, further bolstered by the low numbers of individuals and associated resources required to carry out such screens.

**Table 3.**
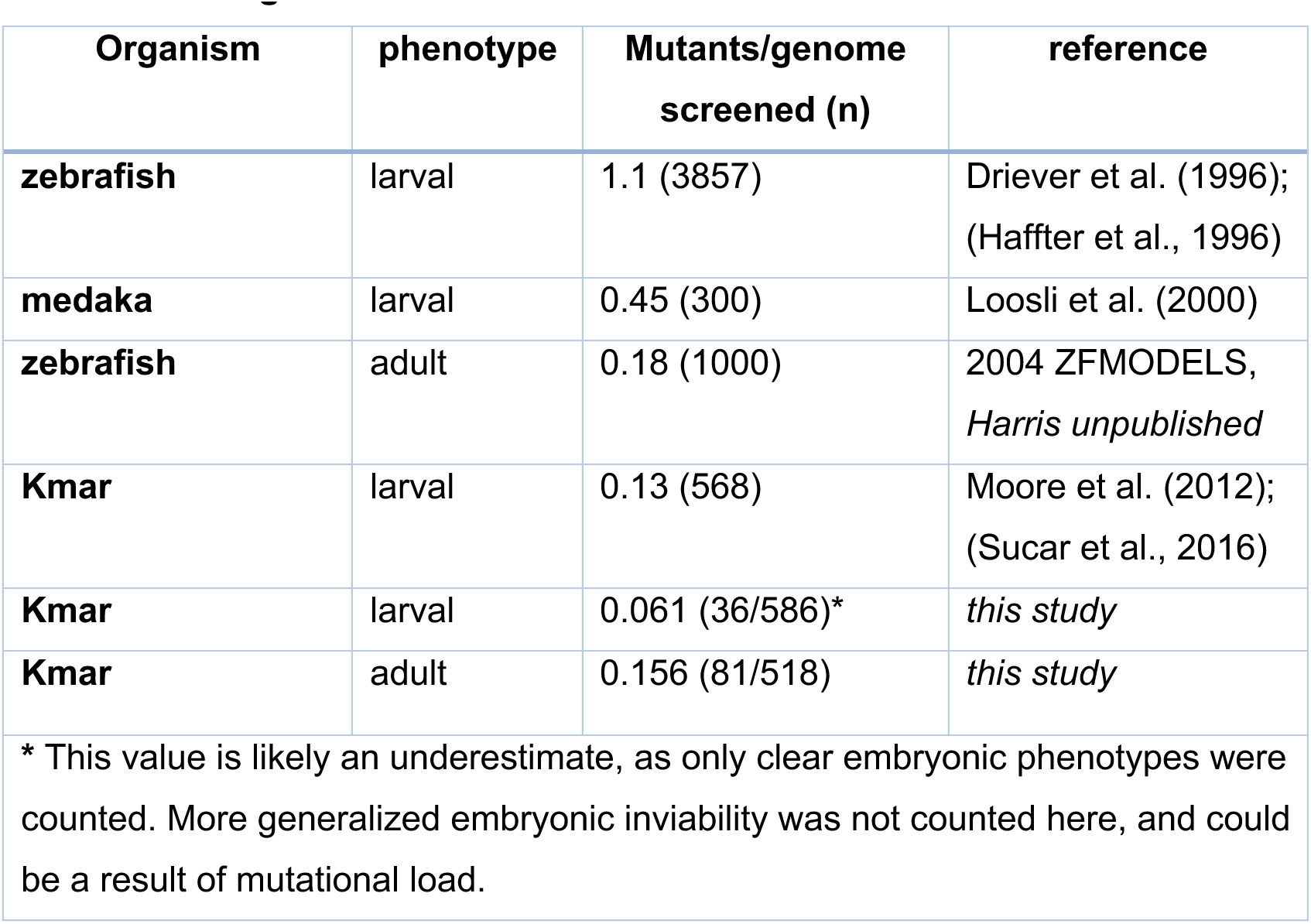
Mutatagenesis in fish models.

| **Organism** | **phenotype** | **Mutants/genome screened (n)** | **reference** |
| --- | --- | --- | --- |
| **zebrafish** | larval | 1.1 (3857) | Driever et al. (1996);<br>(Haffter et al., 1996) |
| **medaka** | larval | 0.45 (300) | Loosli et al. (2000) |
| **zebrafish** | adult | 0.18 (1000) | 2004 ZFMODELS,<br>*Harris unpublished* |
| **Kmar** | larval | 0.13 (568) | Moore et al. (2012);<br>(Sucar et al., 2016) |
| **Kmar** | larval | 0.061 (36/586)* | *this study* |
| **Kmar** | adult | 0.156 (81/518) | *this study* |
| * This value is likely an underestimate, as only clear embryonic phenotypes were counted. More generalized embryonic inviability was not counted here, and could be a result of mutational load. |  |  |  |

A caveat of our screen design is the lack of outcrossing. This presumably leads to maintenance of multiple deleterious alleles in F2 lines, so that some observed phenotypes may be synthetic rather than monogenic. We believe that many of the early phenotypes that showed graded severity of axial defects, eye formation, and cephalic defects were members of this class of complex or compounded phenotypes, as they showed some lineage association with particular founders. In designing future screens, this can be addressed with use of a lower mutagen dose or outcrossing of the first mutagenized stocks, e.g. by mutagenizing only pigmented males, and crossing them to otherwise wild-type albino hermaphrodites. The mutants we maintained were clearly from individual F1 founders and presented with a clean Mendelian ratio in F2 progeny. Mutants were retained with a bias towards useful or informative phenotypes for use in genetic analysis more broadly; all others were not kept.

### Optimizing recovery of diapaused embryos

We further optimized a last hurdle for use of Kmar in genetic analysis. Once eggs are collected, diapaused Kmar larvae frequently fail to hatch. We found that simple addition of pronase (Kanamori et al., 2006) was the most effective treatment and alleviated this experimental barrier. (**Figure 5**). This suggests that diapaused embryos hatch when the chorion becomes permeable or otherwise breached, and that perhaps environmental microbes or other cues that stimulate production of hatching enzyme are often lacking in laboratory cultures. In general support of this, hatching of one embryo in a clutch, which would release hatching enzyme into a small volume of water, often triggers synchronous hatching in others (JAF and ESH, *unpublished observation*). In addition, viable diapaused embryos become active immediately upon mechanical rupture of the chorion.

### Improved egg production supports genetic manipulation

We fortuitously discovered that the *kissylips* mutant has an unusually large egg output. As shown in **Figure 4**, this is true whether eggs are collected frequently or infrequently. We observe that *kissylips* hermpaphrodites are much more likely to deposit eggs in the bottom of their tank than wild-type fish, which strongly prefer to oviposit in the polyester fiber mop. These can remain uneaten for weeks, often resulting in hatchlings sharing a tank with the mother. These observations suggest that *kissylips* hermaphrodites have both unusually frequent oviposition and are less likely to eat eggs once they are laid. The combination of the *kissylips* mutant with frequent removal of eggs overcomes a major barrier to Kmar experimentation, the inability to access suitable numbers of progeny for injections. This set the stage for generation of albino *tyrosinase* mutants with CRISPR/Cas9 nucleases, which are to our knowledge the first genome-edited *K. marmoratus*. From the frequency of albino F1, we can conclude that the founder was a genetic mosaic. This is consistent with our injections, which occurred after the first cleavage. Though early teleost fish blastomeres are often open to the yolk basally, diffusion of editing reagents between blastomeres is apparently not rapid in *K. mamoratus*.

### Plastic traits and generational variation

Many phenotypes obtained in our pilot mutagenesis screen were quite robust, remaining penetrant across generations. However, several lines showed considerable loss of expressivity despite being derived from isogenic, homozygous founders through selfing. This raises the possibility that acquired suppression of some phenotypes may be generated over generations. While we cannot exclude effects of subtle changes in husbandry, these instances harken to findings of epigenetic silencing of traits that can occur in a transgenerational manner (e.g. Bertozzi and Ferguson-Smith, 2020; Chey and Jose, 2022; Verdikt et al., 2023). Given the plasticity of Kmar development to environmental signals, there may be intrinsic mechanisms of trait suppression or modification not seen in traditional model species. Perturbations or challenges (*e.g.* pharmaceutical treatment or environmental shift) could open up this process to be studied using genetic approaches.

## Summary

*Kryptolebias marmoratus* is a truly unique verterbate that invites investigation of dynamic phenotypic states not observed in other common laboratory species. Kmar demographics and variability in life history traits are well documented (Mackiewicz et al., 2006b; Tatarenkov et al., 2012; Tatarenkov et al., 2007; Tatarenkov et al., 2009; Taylor et al., 2008), and plastic responses to environment are beginning to be analyzed (Rossi et al., 2020; Turko and Rossi, 2022). We provide advances in husbandry and genetic tools to enable the integrative exploration development and physiology in the context of a changing environment.

## Acknowledgements

We are grateful to Ryan Earley for assistance in establishing a *K. marmoratus* colony at the University of Maryland, Scott Juntti for his advice and support in performing gene editing experiments, and Ivan Blanco for performing pilot CRISPR experiments.

